# Larva in the loop, a closed loop machine interface system for *Danio rerio* larvae

**DOI:** 10.1101/2024.07.12.603215

**Authors:** John Jutoy, Erica Jung

## Abstract

The optokinetic response (OKR) in Zebrafish (Danio Rerio) had been characterized for its robust response to visual stimuli. Expanding on these works, we developed a novel closed loop control schema to drive a robot utilizing the OKR of Zebrafish larvae. Our system keeps the body of a larva constrained via a novel agarose mold holder that allows for eye movement and vision. The larva is then put under a microscope camera and processed through computer vision to track its eyes via ellipse fitting. Relative eye angle data is then parsed through an algorithm and used to send movement signals to a robot on a lined track. Simultaneously, the robot returns its relative position with respect to the line and converts that information into an OKR stimulation animation which is displayed on an LCD screen in the ventral plane of the larva, thus closing the loop. Through this work we show the capability of larvae OKR to keep a robot on a linear track after an initial oblique entrance to the line. This work displays the potential of our system and how it can pave the way to a Zebrafish Brain-Machine Interface.

## 1. Introduction

To understand and perhaps control the human brain, it is key to have tools that can monitor and elicit responses in the brain. Although great strides have been made in mapping and interfacing with the human brain, there is merit to studying organisms with less complex brain systems such as C. *Elegans, Drosophila, and Danio Rerio*^1^. Animal brain models can help in understanding neuropathology^2^, improving brain mapping^3^, or developing better methodologies for bio-machine interfaces, all of which we hope to address with our system and its future iterations.

Zebrafish larvae are ideal model organisms due to their fast reproduction rate, external development, and translucency^4^. Zebrafish larvae translucency combined with genetic engineering tools and optics (optogenetics), allow for non-invasive neural activity measurement and stimulation^5^. The work presented here describes the progress towards a neural monitoring and elicitation system for Zebrafish larvae.

Thus far, we have been able to develop a visual stimulation system that works in conjunction with a larvae holder. We display the capabilities of the developed Zebrafish Machine Interface which utilizes optokinetic response in both open and closed loop manner. We will discuss how the work presented is a steppingstone towards our goal of a Zebrafish Brain-Machine Interface.

### 1.1 Optokinetic Response

To validate our system, we aimed to reproduce what has been widely done with Zebrafish Larvae which is to demonstrate its optokinetic response through visual stimulation^6^. The Optokinetic response (OKR) is one of the gaze stabilization mechanisms which moves the eye such that a feature of interest is kept in focus.

The Zebrafish larvae OKR has been used for visual screening of mutants due to its robust and reliable response^6^ and has been used to quantify Zebrafish visual acuity^7^. The work presented here capitalizes on this robust response by introducing it in a closed feedback system. We posit that the OKR response of the Zebrafish can maintain a set-point with proper parameter tuning of OKR stimulation.

### 1.2 Novel Larvae Fixation

Consistent and accurate eye tracking requires larva head fixation without obstructing eye movement or vision. Agarose embedding^6,8^ became the standard in Zebrafish larvae fixation due to its transparency and non-toxic attributes. Although agarose embedding has become the standard, it is generally a time-consuming process. We introduce in this work a novel and more efficient method for larvae fixation inspired by Copper et. al^9^.

## 2. Methods

A modular approach was taken to develop the Zebrafish Machine Interface so that modules can be easily added or improved in future work. Brief descriptions are provided for individual modules along with the larvae care methodologies.

The open-loop system, OKR visual stimulation, and eye tracking were used in conjunction to verify that the system could properly elicit OKR in larvae.

The closed-loop system is simply the open-loop system with the addition of the robot car module.

### 2.1 Larvae Fixation

A resin 3D printed positive mold was created with features of a Zebrafish larva as seen in figure 1 c. The dimensions were such that it would constrict larva movement while having enough space for eye movement.

**Figure 1.**
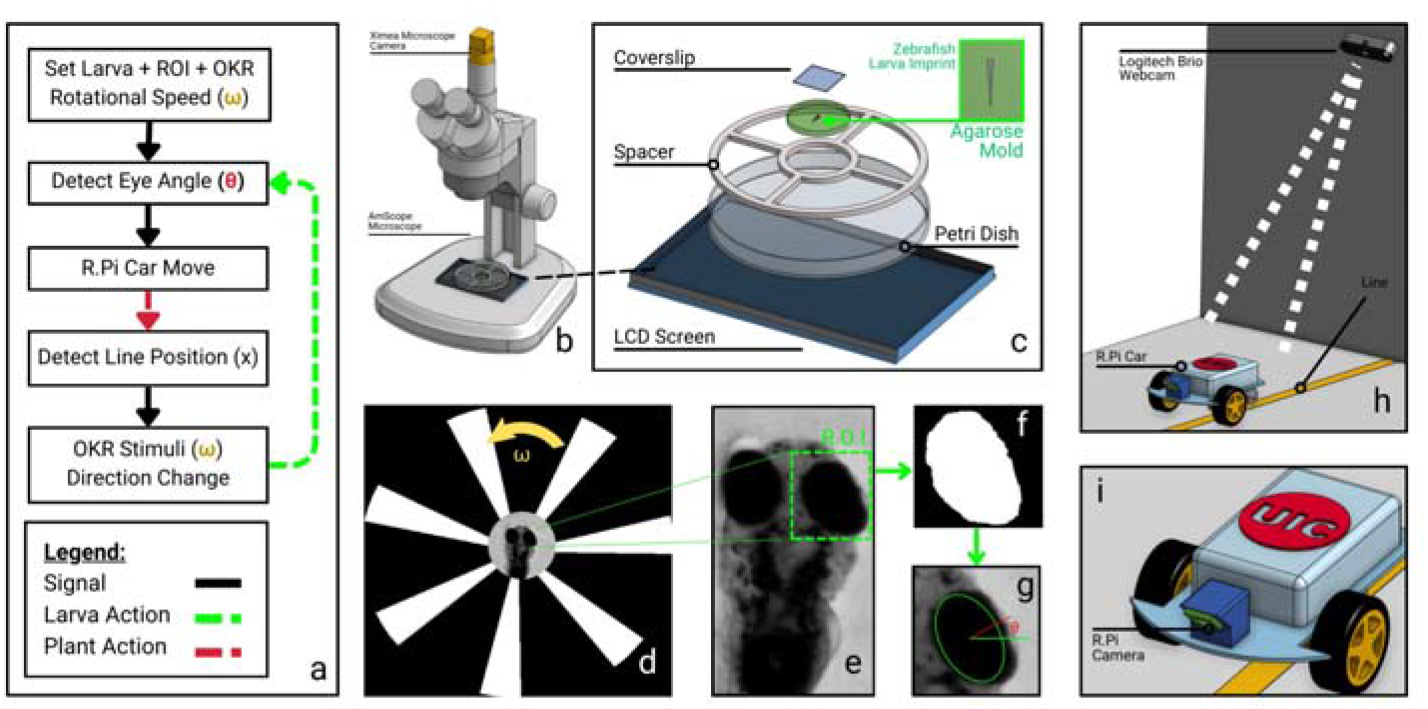
System Setup of Zebrafish Machine Interface including system flow chart, renderings of hardware components, and image processing of eye angle. (a) System logic flowchart with key parameters (ω= constant angular speed of stimuli animation, θ= minor axis angle of fitted ellipse on larva eye with respect to positive horizontal axis, x = distance between center of R.Pi camera and center of yellow line) highlighted and colored. (b) Rendering of Zebrafish system (c) Closeup of Zebrafish larva holder. (d) Animation of rotating black and white arcs at constant angular speed ω provided by LCD screen to elicit optokinetic response. (e) Zoomed in image of greyscale larva video frame along with region of interest (R.O.I.) highlighted. (f) R.O.I. passed through Gaussian filter to isolate larva eye. (g) Ellipse fitted to (f) overlayed into (e) to extract θ. (h) Rendering of R.Pi system. (i) Zoomed in rendering of R.Pi car to display the horizontal distance between camera and line.

**Figure 2.**
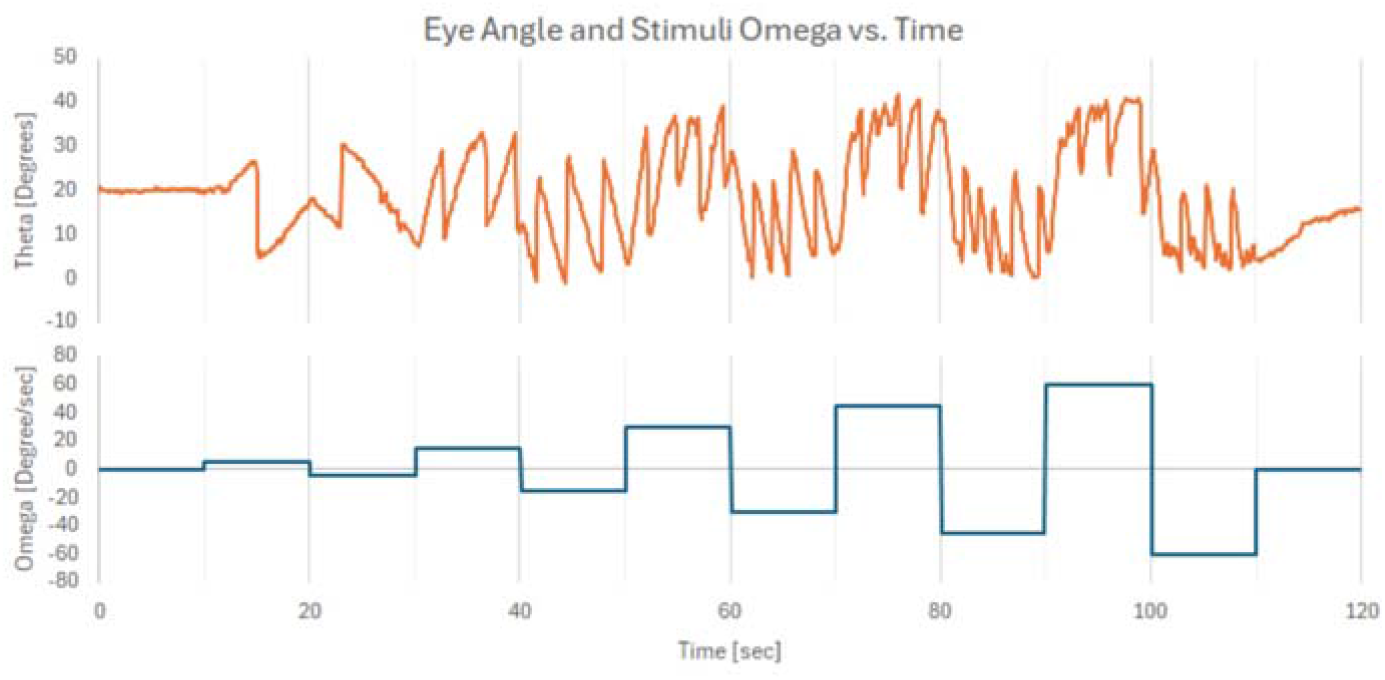
Right eye angle response for a single Zebrafish larva at varying direction and ω as seen in figure 1-d. Eye angles are in terms of degrees while ω are in terms of degrees/sec. Note that eye angle slope direction agrees with stimuli ω direction within times that the stimuli was given: 20-120 sec.

Creation of the device consisted of heating up agarose gel into its liquid form and placing it into a petri dish with the positive mold. The agarose solidifies into a translucent gel in which the positive mold can be carefully removed. Negative space left by the positive mold is then filled with a larva through a hair loop device ^10^.

### 2.2 OKR Visual Stimulation

A module in the system software was created to provide a rotating grating animation using the OpenCV python library. Several parameters can be easily modified by the system including grating rotation speed *ω*, number of gratings, grating arc length, and grating radius. Sufficient contrast for the eye tracking module was provided by including a blank white circle directly under the fish.

The parameters chosen for testing were identified manually based on visual inspection of OKR from the larvae. These parameters were found to be: 5 gratings, 50% grating spacing (ie: half of the grating is black, and half is white).

### 2.3 Eye Tracking

The fish eye tracking module converts images from the microscope camera video stream into eye angles of the Zebrafish eye of interest. Using the OpenCV library of python, raw images are converted to grey scale, blurred, then threshold filtered to a value found suitable for isolation of the Zebrafish eye. Contouring is done on the isolated eye and an ellipse is then fitted to it. The major axis of the ellipse is then used as the angle of the eye.

### 2.4 Robot Car

A raspberry pi car with two wheeled motors in the front and a free moving wheel in the back was chosen as the vehicle for the Zebrafish to control. The modified chassis was made to accommodate a raspberry pi camera. Communication to the raspberry pi car from the main workstation was done through the Paramiko library.

The robot car was tracked through a Logitech Brio webcam and processed through OpenCV to extract its estimated position.

### 2.5 Zebrafish Larvae

Larvae were bred from wild-type adult in a dedicated fish room, temperature controlled to 26^⍰^*C*, with a 14-h light and 10-h darkness cycle. The larvae are harvested and kept in the fish room until 4-5 days post fertilization (dpf) from which they are brought to the testing facility a day prior for experiments. Larvae are kept in an incubation chamber set at 27^⍰^*C* outside of experimental trials.

## 3. Results

### 3.1 Verification of Optokinetic Response

To verify that OKR is displayed by Zebrafish larvae, a simple test with a duration of 140 seconds was created to analyze eye response to the developed OKR stimuli. The first and last 10 seconds of the test hide the individual arc features of the stimuli to serve as a baseline of no response and is referred to as *ω* = 0. The stimuli are displayed at *ω* = 5,15,30,45,60 degrees / sec in 20 second time intervals between 20 - 120 respectively. Within each 20 second time bin, a direction switch is made.

Analysis of total eye angle response (figure 3) was done to display that the OKR response was related to the visual stimuli of the system.

**Figure 3.**
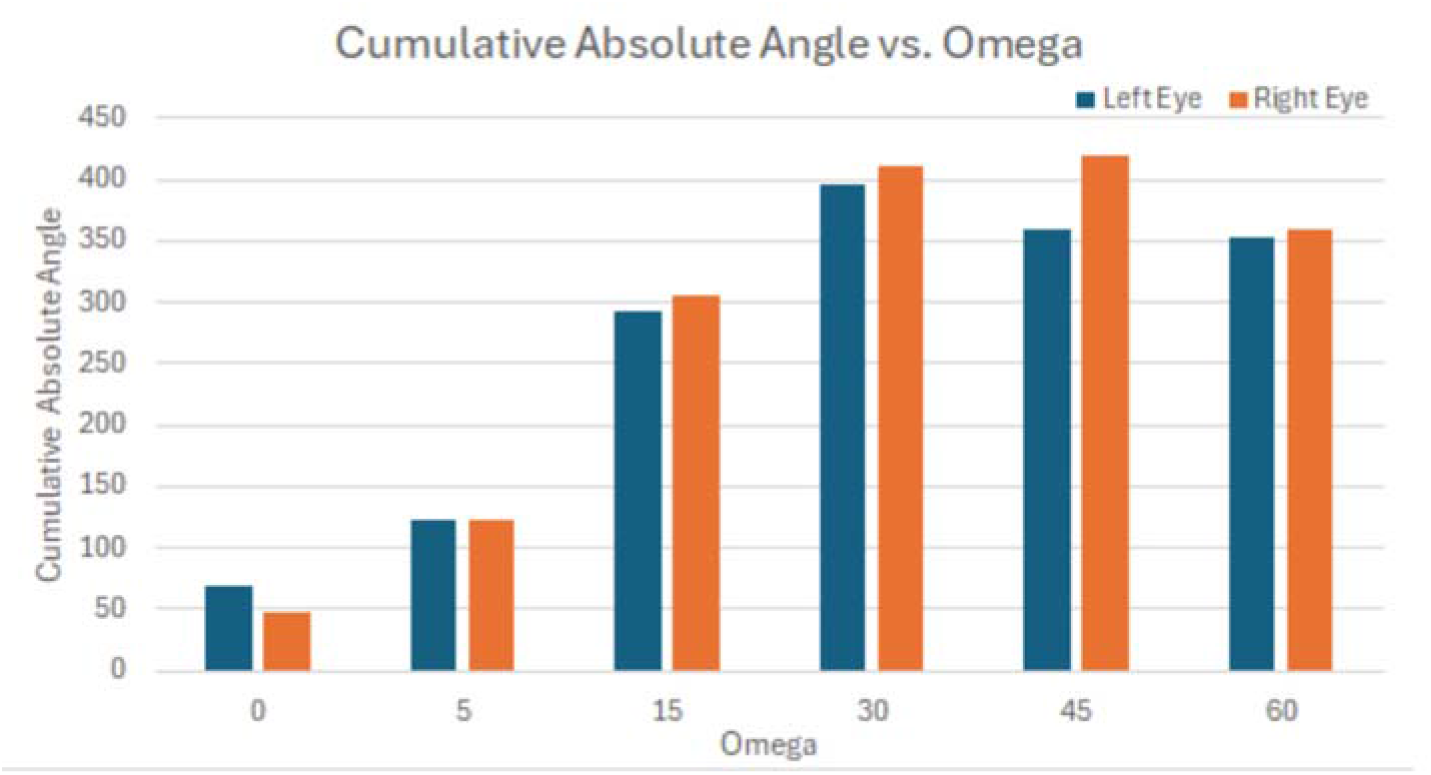
Absolute cumulative eye angle of left and right eye compared to stimuli speed ω was done to display that larvae response is due to our visual stimuli system. ω = 0 represents no visual stimuli present.

**Figure 4.**
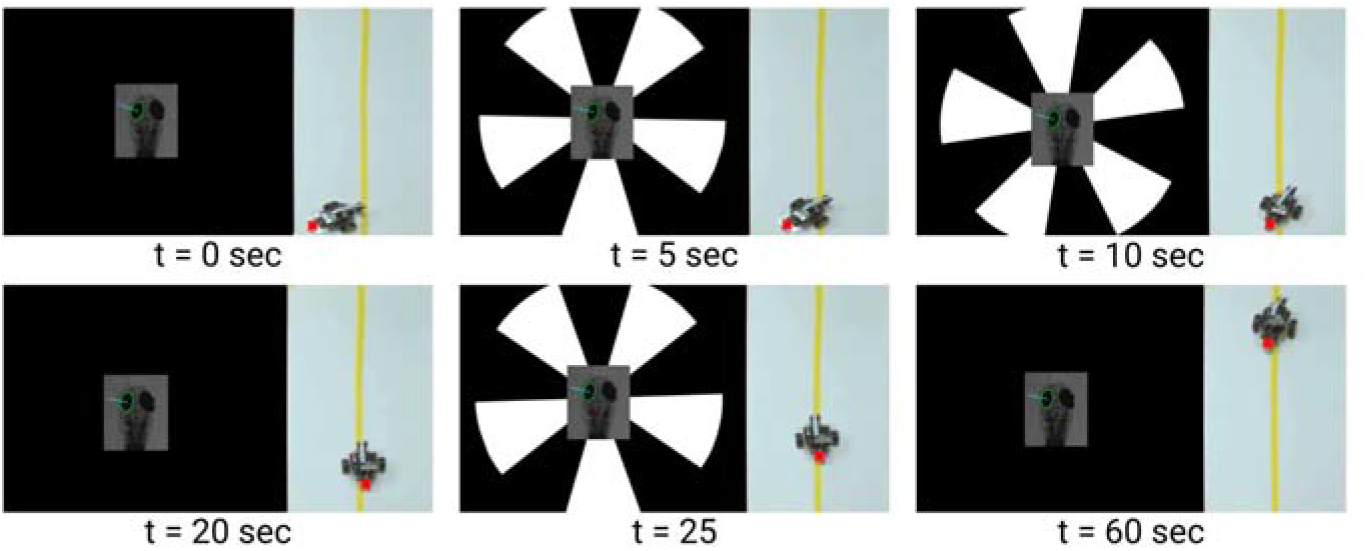
Larvae in the loop line following real time view. A larva with its left eye being tracked superimposed on top of the visual stimuli its receiving (left) and the robot car being controlled by the Zebrafish larva (right). The change in left eye angle is checked with a threshold in order to send a move signal (left if |θ| < threshold, right if |θ| > threshold, and forward if within threshold) A camera in front of the robot car sends the location of where it last saw the yellow line to the visual stimuli system adjusting direction/display respectively in order to direct the larva’s eye to back to the direction of the yellow line. Note that ω is kept constant throughout the driving trial.

### 3.2 Larvae in the Loop Line Following

Once optokinetic response was identified in a fish, trials were conducted to see if the larvae can maintain a robot car within a line at different *ω* as seen in figure 5. A representative set of trials from a single larva driver can be seen in figure 6. Each trial consisted of starting the robot car facing the yellow line at a slight random oblique angle. The car was tracked and driven by the larvae for a duration of 45 seconds.

**Figure 5.**
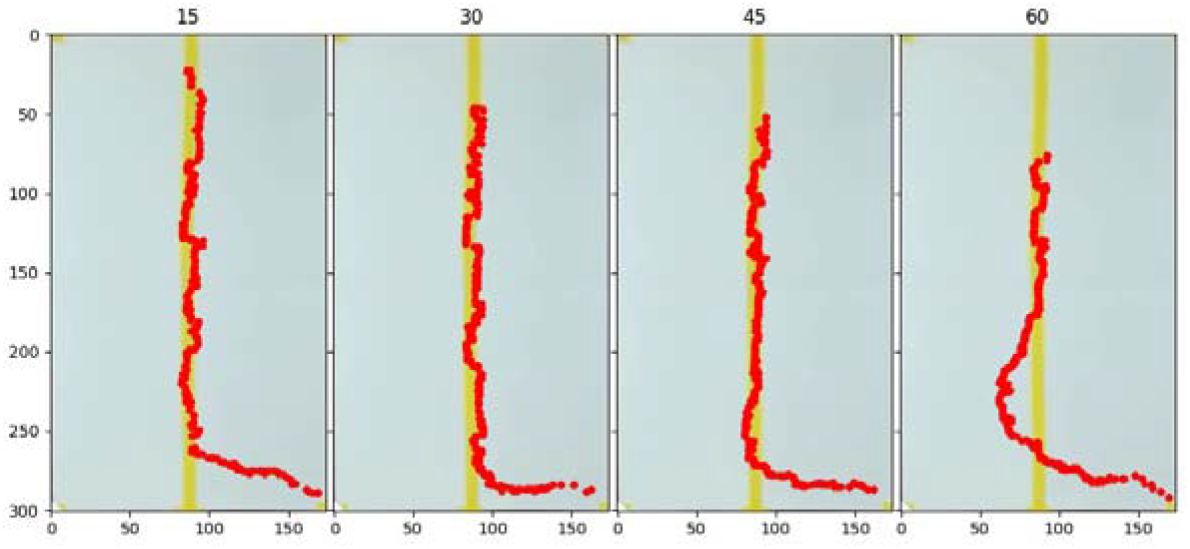
Pathway traces of the Zebrafish driven car at different ω. The top number on the plots represents ω in degrees / sec and each plot represents a trial with a duration of 45 seconds. Axis is in terms of pixels. The robot car is started initially at the bottom right.

## 4. Discussion

### 4.1 Novel Larvae Fixation Method

The larvae fixation method introduced in this work succeeded in keeping larvae in place to run cumulative visual experiments up to several minutes long. Visual experiments can elicit startle responses, which were observed in some trials, still our device was able to keep the larvae fixed and within the region of interest. For the use case in this paper, this device made trials significantly more efficient as minor decoupling of a larvae and the device can be tolerated but may be inappropriate for other cases where slight movements are not tolerable.

### 4.2 Mechanism of Visual Stimuli

The mechanism behind optokinetic stimuli projected from below the larvae was not the focus of this project. We posit that it may be due to the fish responding to the visual stimuli below it as in optomotor response setups. Alternatively, it may be due to the refraction of light across various interface mediums that are causing a frontal-lateral projection to the larvae. It may also be a combination of the two mentioned theories. Regardless, optokinetic response is clearly present from the projection of the stimuli below the larvae as seen in 2 as the eye angle response agrees with the stimuli direction and is sporadic or not present when the stimuli is either not displayed or paused.

### 4.3 Movement Thresholds

Individual larvae display variation in OKR sensitivity. As such, the movement threshold utilized for an OKR sensitive larva was not sufficient for a larva with less OKR sensitivity. Future work should identify this threshold and compare the driving of larvae with varying OKR sensitivity. Variation in OKR sensitivity may be due to the below ventral projection of visual stimuli as discussed above. Another explanation may be due to slight misalignments between larva, fixation device, and microscope. Since there is a range of tolerances depending on larvae size and hydration of the agarose, slight dorsal misalignments were noticed in some trials. However, there is also possibility that these OKR sensitivities observed through this system may be an aspect of the Zebrafish larvae not investigated before.

### 4.4 Future Work

We emphasize that the work presented here is a fundamental step towards developing a Zebrafish Brain-Machine Interface. To map the brain of an organism through high throughput inputs, we require a way to simultaneously stimulate and record the larvae. In addition to this, we must keep the larvae in place without interfering with the input stimuli and output responses. Although the current output responses we utilize in this work are not explicitly neural signals, we demonstrate the potential of our system to visually stimulate the larvae which we aim to map to neural activity.

Further characterization besides *ω* will be explored in the future. Visual parameters such as the spacing between gratings, number of gratings, and distance of gratings from the larvae may affect the response. As mentioned in the OKR section of methodology, the visual parameters were tuned manually based on reactivity of the specific larvae. Whether there exist relationships between these parameters and the larvae will be analyzed in detail in a future paper.

Work is planned to further explore the novel larvae fixation method described in this short paper. Although the overarching concept was provided, in depth discussion was not included here as current work is ongoing to characterize fixation parameters.

## 5. Conclusion

This work displays that Zebrafish larvae can be utilized as controllers to simple systems like line following robots. Further experimentation is required to elucidate system relationships, however, with coarse parameter settings (i.e. arbitrary setting of thresholds) promising results can be gained. Furthermore, we briefly showcased our novel, fast, and efficient technique for fixing a larva in place with the trade-off that startled larvae may get slightly misaligned.

## Acknowledgments

This work was supported by the National Science Foundation [Neuron-to-Neuron Interface: Optically Connected Neurons Between the Brains of Two Zebrafish. Grant#: 2309589].

